# Through-plane super-resolution with autoencoders in diffusion magnetic resonance imaging of the developing human brain

**DOI:** 10.1101/2021.12.06.471406

**Authors:** Hamza Kebiri, Erick J. Canales Rodríguez, Hélène Lajous, Priscille de Dumast, Gabriel Girard, Yasser Alemán-Gómez, Mériam Koob, András Jakab, Meritxell Bach Cuadra

## Abstract

Fetal brain diffusion magnetic resonance images are often acquired with a lower through-plane than in-plane resolution. This anisotropy is often overcome by classical upsampling methods such as linear or cubic interpolation. In this work, we employ an unsupervised learning algorithm using an autoencoder neural network to enhance the through-plane resolution by leveraging a large amount of data. Our framework, which can also be used for slice outliers replacement, overperformed conventional interpolations quantitatively and qualitatively on pre-term newborns of the developing Human Connectome Project. The evaluation was performed on both the original diffusion-weighted signal and on the estimated diffusion tensor maps. A byproduct of our autoencoder was its ability to act as a denoiser. The network was able to generalize to fetal data with different levels of motion and we qualitatively showed its consistency, hence supporting the relevance of pre-term datasets to improve the processing of fetal brain images.

## 1 INTRODUCTION

The formation and maturation of the white matter is at its highest rate during the fetal stage of human brain development. To have more insight into this critical period, *in utero* brain imaging techniques offer a unique opportunity. Diffusion weighted magnetic resonance imaging (DW-MRI) is a well-established tool to reconstruct in vivo and non-invasively the white matter tracts in the brain (Basser and Pierpaoli, 2011; Johansen-Berg and Behrens, 2013). Fetal DW-MRI, in particular, could characterize early developmental trajectories in brain connections and microstructure (Bui et al., 2006; Huang et al., 2009; Huang and Vasung, 2014; Jakab et al., 2015). Hence, fetal DW-MRI has been of significant interest for the past years where studies (Righini et al., 2003; Kasprian et al., 2008; Khan et al., 2019) have provided analysis using diffusion tensor imaging (DTI) by computing diffusion scalar maps - such as fractional anisotropy (FA) or mean diffusivity (MD) - using a limited number of gradient directions. A recent work focused on reconstructing fiber Orientation Distribution Functions (fODF) (Deprez et al., 2019) using higher quality datasets and rich information including several gradient directions (32 and 80), and higher b-values (750 and 1000 *s/mm*^2^) and signal-to-noise ratio (SNR) (3 Tesla magnetic field strength). Additionally, the datasets were acquired in a controlled and uniform research setting with healthy volunteers, which can hardly be reproduced in the clinical environment.

Albeit promising results, acquiring high quality data remains a main obstacle in the field of fetal brain imaging. Firstly, unpredictable and uncontrollable fetal motion is a major challenge. To overcome this problem, fast echo-planar imaging (EPI) sequences are typically used to freeze intra-slice motion. However, intra- and inter-volume motion still has to be addressed in the post-processing steps using sophisticated slice-to-volume registration (SVR) (Kim et al., 2010; Fogtmann et al., 2013; Marami et al., 2016). Moreover, EPI sequences generate severe non-linear distortions that need adapted distortion correction algorithms (Kuklisova-Murgasova et al., 2017). Additionally, the resulting images display low SNR due to at least three factors: the inherently small size of the fetal brain, the surrounding maternal structures and amniotic fluid, and the increased distance to the coils. In order to compensate for the low SNR in EPI sequences, series with thick voxels (i.e. low through-plane resolution) are often acquired. Finally, to shorten the acquisition time, small b-values (*b* = 400 – 700*s/mm*^2^) and a low number of gradient directions (10-15) (Kasprian et al., 2008; Khan et al., 2019) are commonly used in fetal imaging, which in turn will result in a low angular resolution.

Clinical protocols typically acquire several anisotropic orthogonal series of 2D thick slices to cope with high motion and low SNR. Then, super-resolution reconstruction techniques that have been originally developed for structural T2-weighted images (Rousseau et al., 2006; Gholipour et al., 2010; Kuklisova-Murgasova et al., 2012; Tourbier et al., 2015; Kainz et al., 2015; Ebner et al., 2020) by combining different 3D low resolution volumes, have also been successfully applied in 4D fetal functional (Taymourtash et al., 2021) and diffusion MRI contexts (Fogtmann et al., 2013; Deprez et al., 2019). Still, despite these two pioneer works, super-resolution DW-MRI from multiple volumes has been barely explored in vivo. In fact, the limited scanning time to minimize maternal discomfort hampers the acquisition of several orthogonal series, resulting in a trade-off between the number of gradient directions and orthogonal series. Thus, DW-MRI fetal brain protocols are not standardized from one center to another (Table S1 in Supplementary Material) and more experiments have to be conducted in this area to design optimal sequences (Kebiri et al., 2021a,b).

Nevertheless, fetal DW-MRI resolution enhancement could also benefit from *single image superresolution* approaches, i.e. either within each DW-MRI 3D volume separately or using the whole 4D volume including all diffusion measurements. In fact, it has been demonstrated that a linear or cubic interpolation of the raw signal enhances the resulting scalar maps and tractography (Dyrby et al., 2014). In practice, this is typically performed either at the signal level or at DTI scalar maps (Jakab et al., 2017). We believe that single volume and multiple volumes super-resolution can also be performed together, i.e. where the output of the former is given as the input of the later. This aggregation could potentially lead to a better motion correction and hence to a more accurate final high resolution volume.

Several studies have proposed single image super-resolution enhancement methods for DW-MRI but, to the best of our knowledge, none of them has been applied neither to anisotropic datasets nor to the developing brain. In Coupé et al. (2013), the authors utilized a non-local patch-based approach in an inverse problem paradigm to improve the resolution of adult brain DW-MRI volumes using a non diffusion weighted image (*b* = 0*s/mm*^2^) prior. Although this approach yielded competitive results, it was built upon a sophisticated pipeline which made it not extensively used. The first machine learning study (Alexander et al., 2017) have used shallow learning algorithms to learn the mapping between diffusion tensor maps of a downsampled high resolution image and the maps of the original image. Recently, deep learning models which can implicitly learn relevant features from training data were used to perform single image super-resolution with a convolutional neural network (Elsaid and Wu, 2019; Dong et al., 2015) and a customized U-Net (Chatterjee et al., 2021; Ronneberger et al., 2015). Both approaches produced promising results in a *supervised* learning scheme. Supervision needs however large high quality datasets that are scarce for the perinatal brain for the reasons enumerated above.

The specific challenge of fetal DW-MRI is 3-5 mm acquired slice thickness, with only a few repetition available. Hence our main objective is to focus on through-plane DW-MRI resolution enhancement. This would be valuable not only for native anisotropic volumes but also for outlier slice recovery. In fact, motion-corrupted slices in DW-MRI are either discarded, which results in a loss of information, or replaced using interpolation (Niethammer et al., 2007; Chang et al., 2005; Andersson et al., 2016). We approached this problem from an image synthesis point of view using *unsupervised* learning networks such as autoencoders (AEs), as demonstrated in cardiac T2-weighted MRI (Sander et al., 2021) and recent works in DW-MRI (Chung et al., 2021). Here we present a framework with autoencoders that are neural networks learning in an *unsupervised* way to encode efficient data representations and can behave as generative models if this representation is structured enough. By accurately encoding DW-MRI slices in a low-dimensional latent space, we were able to successfully generate new slices that accurately correspond to in-between “missing” slices. In contrast to the above referred *supervised* learning approaches, this method is scale agnostic, i.e. the enhancement scale factor can be set *a posteriori* to the network training.

Realistically increasing the through-plane resolution would potentially help the clinicians to better assess whether the anterior and posterior commissures are present in cases with complete agenesis of the corpus callosum (Jakab et al., 2015). It can reduce partial volume effects and thus contribute to the depiction of more accurate white matter properties in the developing brain.

In this work, we present the first unsupervised through-plane resolution enhancement for perinatal brain DW-MRI. We leverage the high quality dataset of the developing Human Connectome Project (dHCP) where we train and quantitatively validate on pre-term newborns that are anatomically close to fetal subjects. We finally demonstrate the performance of our approach in fetal brains.

## 2 MATERIALS AND METHODS

### 2.1 Data

#### 2.1.1 Pre-term dHCP data

We selected all the 31 pre-term newborns of 37 gestational weeks (GW) or less at the time of scan (range: [29.3,37.0], mean: 35.5, median: 35.7) from the dHCP dataset (Bastiani et al., 2019) (subject IDs in Table S2 of Supplementary Material). Acquisitions were performed using a 3T Philips Achieva scanner (32-channel neonatal head-coil) with a monopolar spin-echo EPI Stejksal-Tanner sequence (Δ=42.5ms, *δ*=14ms, TR=3800ms, TE=90ms). The spatial resolution was 1.17×1.17×1.5 *mm*^3^ with a field of view of 128×128×64 voxels. The dataset was acquired with a multi-shell sequence using four b-values (b ∈ {0, 400, 1000, 2600}*s/mm*^2^) with 300 volumes but we have only extracted the 88 volumes corresponding to b = 1000*s/mm*^2^ (b1000) as a compromise of high contrast-to-noise ratio (CNR), i.e. b1000 has a higher CNR than b400 and b2600 (Bastiani et al., 2019), and proximity to the *b* = 700*s/mm*^2^ that is typically used in clinical settings for fetal DW-MRI. Brain masks and region/tissue labels segmented using a pipeline based on Draw-EM algorithm (Makropoulos et al., 2018, 2014) were available in the corresponding anatomical dataset. All the images were already corrected for inter-slice motion and distortion (susceptibility, eddy currents and motion).

#### 2.1.2 Fetal data

Fetal acquisitions were performed at 1.5T (MR450, GE Healthcare, Milwaukee, WI, USA) in the University Children’s Hospital Zürich (KISPI) using a single-shot EPI sequence (TE=63 ms, TR=2200 ms) at *b* = 700*s/mm*^2^ (b700). The acquisition time was approximately 1.3 minutes per 4D volume. The in-plane resolution was 1×1 *mm*^2^, the slice thickness 4-5*mm* and the field of view 256×256×14 – 22 voxels. Three axial series and a coronal one were acquired for each subject. Brain masks were manually generated for the b0 (*b* = 0*s/mm*^2^) of each acquisition and automatically propagated to the diffusion-weighted volumes. Between 8 and 18 T2-weighted images were also acquired for each subject where corresponding brain masks were automatically generated using an in-house deep learning based method using transfer learning from Salehi et al. (2018). Manual refinements were needed for a few cases at the brain boundaries.

#### 2.1.3 Fetal data processing

We selected three subjects with high quality imaging and without motion artefacts (24, 29 and 35 GW) and three subjects with a varying degree of motion (23, 24 and 27 GW). Figure 1 illustrates a DW-MRI volume of a motion-free case (29 GW). By performing quality control, we discarded highly corrupted volumes due to motion resulting in severe signal drops in two moving subjects and very low SNR volumes in one motion-free subject (Table S3 in Supplementary Material shows the discarded volumes for each subject). All the subjects were pre-processed for noise, bias field inhomogeneities and distortions using the Nipype framework (Gorgolewski et al., 2011). The denoising was performed using a Principal Component Analysis based method (Veraart et al., 2016), followed by an N4 bias-field inhomogeneity correction (Tustison et al., 2010). Distortion was corrected using an in-house implementation of a state-of-the-art algorithm for fetal brain (Kuklisova-Murgasova et al., 2017) consisting in rigid registration (Avants et al., 2009) of a structural T2-weighted image to the b0 image, followed by a non-linear registration (Avants et al., 2009) in the phase-encoding direction of the b0 to the same T2-weighted image. The transformation was then applied to the diffusion-weighted volumes. A block matching algorithm for symmetric global registration was also performed for two subjects with motion (NiftyReg, Modat et al., 2014). The b0 image of the first axial series was selected as a reference to which we subsequently registered the remaining volumes, i.e. the non b0 images from the first axial and all volumes from the two others. Gradient directions were rotated accordingly.

**Figure 1.**
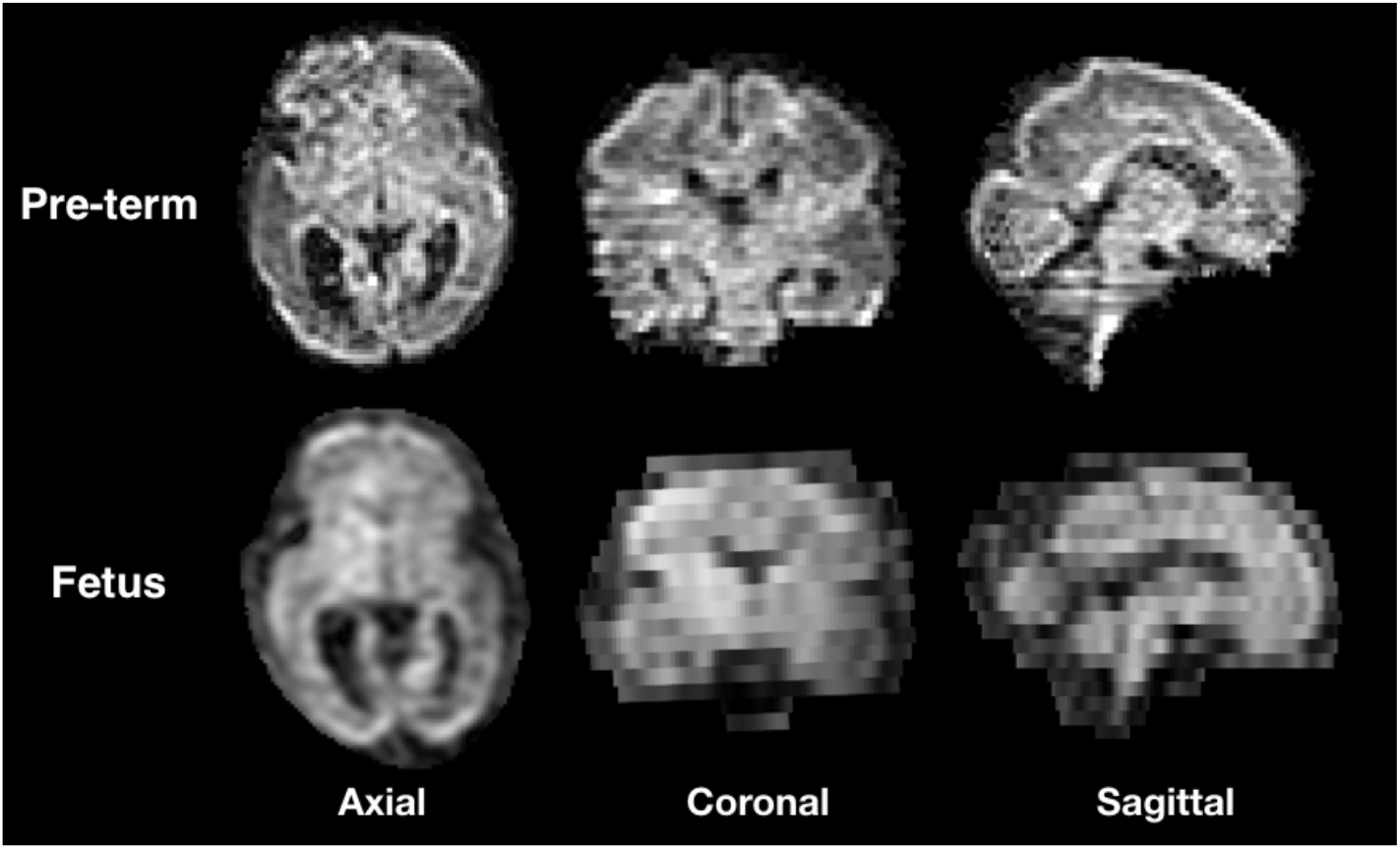
Illustration of the three orientations of a DW-MRI volume from a still fetal subject (29 GW) and pre-term newborn of the same gestational age.

### 2.2 Model

#### 2.2.1 Architecture

Our network architecture, similarly to Sander et al. (2021), is composed of four blocks in the encoder and four in the decoder (Figure 2). Each block in the encoder consists of two layers made of 3×3 convolutions followed by a batch normalization (Ioffe and Szegedy, 2015) and an Exponential Linear Unit non-linearity.

**Figure 2.**
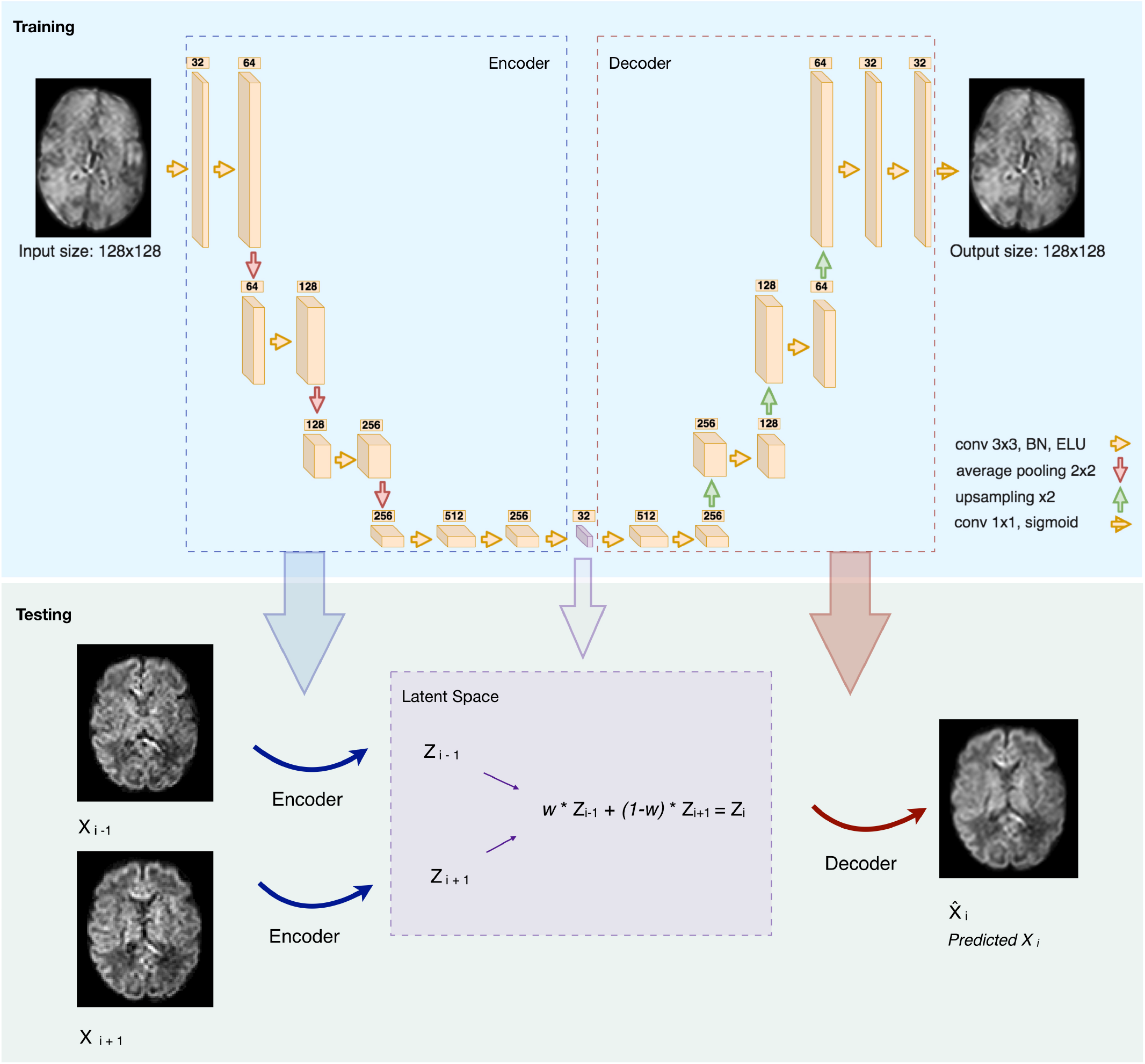
Illustration of the network architecture (top): Each box is a multi-channel feature maps. The number of channels is denoted on top of each box. The violet box represents the latent space of the autoencoder (BN: Batch Normalization. ELU: Exponential Linear Unit). An illustration of how we generated middle slice(s) is shown in the bottom panel (Testing), for the case of an equal slice weighting (w=0.5) and b1000.

The number of feature maps is doubled from 32 after each layer and the resulting feature maps are average-pooled. We further added two layers of two 3×3 convolutions in which the feature maps of the last layer was used as the latent-space of the autoencoder. The decoder uses the same architecture as the encoder but by conversely halving the number of feature maps and upsampling after each block using nearest neighbor interpolation. At the final layer a 1×1 convolution using sigmoid function is applied to output the predicted image. The number of network parameters is 6,098,689.

#### 2.2.2 Training and optimization

In the TensorFlow framework (Abadi et al., 2016), we have trained our network solely on b0 images, using an 8-fold nested cross validation where we trained and validated on 27 subjects and tested on four. The proportion of the validation data was set to 15% of the training set. The training/validation set contains 25,920 slices of a 128×128 field of view, totalling 424,673,280 voxels. Our network was trained in an unsupervised manner by feeding normalized 2D axial slices that are encoded as feature maps in the latent space. The number of feature maps, and hence the dimensionality of the latent space, was optimized (optimal value to 32) using Keras-Tuner (Chollet et al., 2015). The batch size and the learning rate were additionally optimized and set to 32 and 5e-5, respectively. The network was trained for 200 epochs using Adam optimizer (Kingma and Ba, 2014) with the default parameters *β*_1_ = 0.5, *β*_2_ = 0.999, and the network corresponding to the epoch with the minimal validation loss was then selected. Network code and checkpoint example can be found in our Github repository ^1^.

#### 2.2.3 Inference

The network trained on b0 images was used for the inference of b0 and b1000 volumes. Two slices were encoded in the latent space and their N ”in-between” slice(s) (N=1 ,2 in our experiments) were predicted using weighted averages of the latent codes of the two slices. An example on pre-term b1000 data for a weight of 0.5 is shown in Figure 2 (Testing). Similarly, the same b0 network was also used to increase the through-plane resolution of fetal b0 and b700 volumes. Finally, since the network outputs were normalized between 0 and 1, histogram normalization to the weighted average of the input images was performed.

### 2.3 Experiments and evaluation

#### 2.3.1 Pre-term newborns

Our network was separately tested on b0 images and the 88 volumes of b1000 using an 8-fold cross validation where seven folds contain four subjects and one fold contains three subjects. We removed N intermediate slices (N=[1, 2]) from the testing set volumes in an alternating order and used the (weighted) average latent space feature maps of the to-be adjacent slices to encode the N missing slice(s) using the autoencoder (Figure 2, Testing). The resulting latent representation was then decoded to predict the N slices in the voxel space, which were compared to the previously removed N slices, i.e. the ground truth (GT). The same N slices were also generated using three baseline approaches: trilinear, tricubic and B-spline of 5*^th^* order interpolations (using Tournier et al. (2012); Avants et al. (2009)) for comparison. We denote them respectively for removing one or two slices: Linear-1, Cubic-1, Spline −1 and Linear-2, Cubic-2, Spline-2.

##### Latent space validation

We have compared the latent space representations between different gradient directions of all possible pairs from the 88 volumes of the b1000 4D volume.

##### Robustness to noise

We have added different low levels of Rician noise (Gudbjartsson and Patz, 1995) to the original signal as follows: for each pixel with a current intensity *S_clean_*, the new intensity 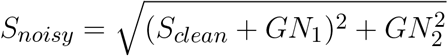 where *GN*_1_ and *GN*_2_ are random numbers sampled from a Gaussian distribution with zero mean and a standard deviation of *S_clean_*(*b* = 0)/*SNR_out_*, and *SNR_out_* is the desired SNR we aim to simulate. Three SNR of {27, 25, 23} and {20, 16, 13} were simulated for b0 and b1000 respectively. We have used higher noise levels for b1000 to better simulate the inherently lower SNR in this configuration.

##### Scalar maps

By merging the b0 and b1000 using the autoencoder enhancement, we reconstructed fractional anisotropy (FA), mean diffusivity (MD), axial diffusivity (AD) and radial diffusivity (RD) from DTI using Dipy (Garyfallidis et al., 2014) separately for AE-1 or AE-2, i.e. where we respectively remove one (N=1) or two slices (N=2). We further subdivided the computation in specific brain regions (cortical gray matter, white matter, corpus callosum and brainstem as provided by the dHCP). Region labels were upsampled and manually refined to match the super-resoluted/interpolated volumes. We performed similar computation of the diffusion maps generated using the trilinear, tricubic and B-spline interpolated signals.

#### 2.3.2 Fetal

For each subject and each 3D volume (b0 or DW-MRI), we generated one or two middle slices using autoencoder enhancement, hence increasing the physical resolution from 1×1×4-5*mm*^3^ to 1×1×2-2.5*mm*^3^ and 1×1×1.33-1.67*mm*^3^ respectively. We then generated whole-brain DTI maps (FA, MD, AD and RD) and show the colored FA. Splenium and genu structures of the corpus callosum were additionally segmented on FA maps for subjects in which these structures were visible. The mean FA and MD were reported for these regions for original and autoencoder enhanced volumes.

#### 2.3.3 Quantitative evaluation

##### Raw diffusion signal

We computed the voxel-wise error between the raw signal synthesized by the autoencoder and the GT using the mean squared error (MSE) and the peak signal-to-noise ratio (PSNR). We compared the autoencoder performance with the three baseline approaches: trilinear, tricubic and B-spline of 5*^th^* order interpolations.

##### Latent space validation

We have computed the average squared Euclidian voxel-wise distance between slices of all 3D b1000 volume pairs. This was performed both at the input space and at the latent space representation.

##### Robustness to noise

We computed with respect to the GT signal the error of the signal with noise, and the output of the autoencoder using the signal with noise as input. We compared the results using MSE separately for b0 and b1000.

##### Scalar maps

We computed the voxel-wise error between the diffusion tensor maps reconstructed with the GT and the one by merging the b0 and b1000 using the autoencoder enhancement. We computed the error separately using either AE-1 or AE-2. We used the MSE and the PSNR as metrics and the same diffusion maps generated using the trilinear, tricubic and B-spline interpolated signal as a baseline. Moreover, we qualitatively compare colored FA generated using the best baseline method, autoencoder and the GT.

## 3 RESULTS

### 3.1 Pre-term newborns

First, we inspected the latent space and how the 88 DW-MRI volumes are encoded with respect to each other. We can notice in Figure 3 (right panel) that as two b-vectors’ angle approaches orthogonality (90°) the difference between the latent representations of their corresponding volumes increases. Conversely the difference decreases the more the angle tends towards 0° or 180°. Although the pattern is more pronounced in the input space (Figure 3, left panel), this trend is a fulfilled necessary condition to the generation of coherent representations of the input data by our network.

**Figure 3.**
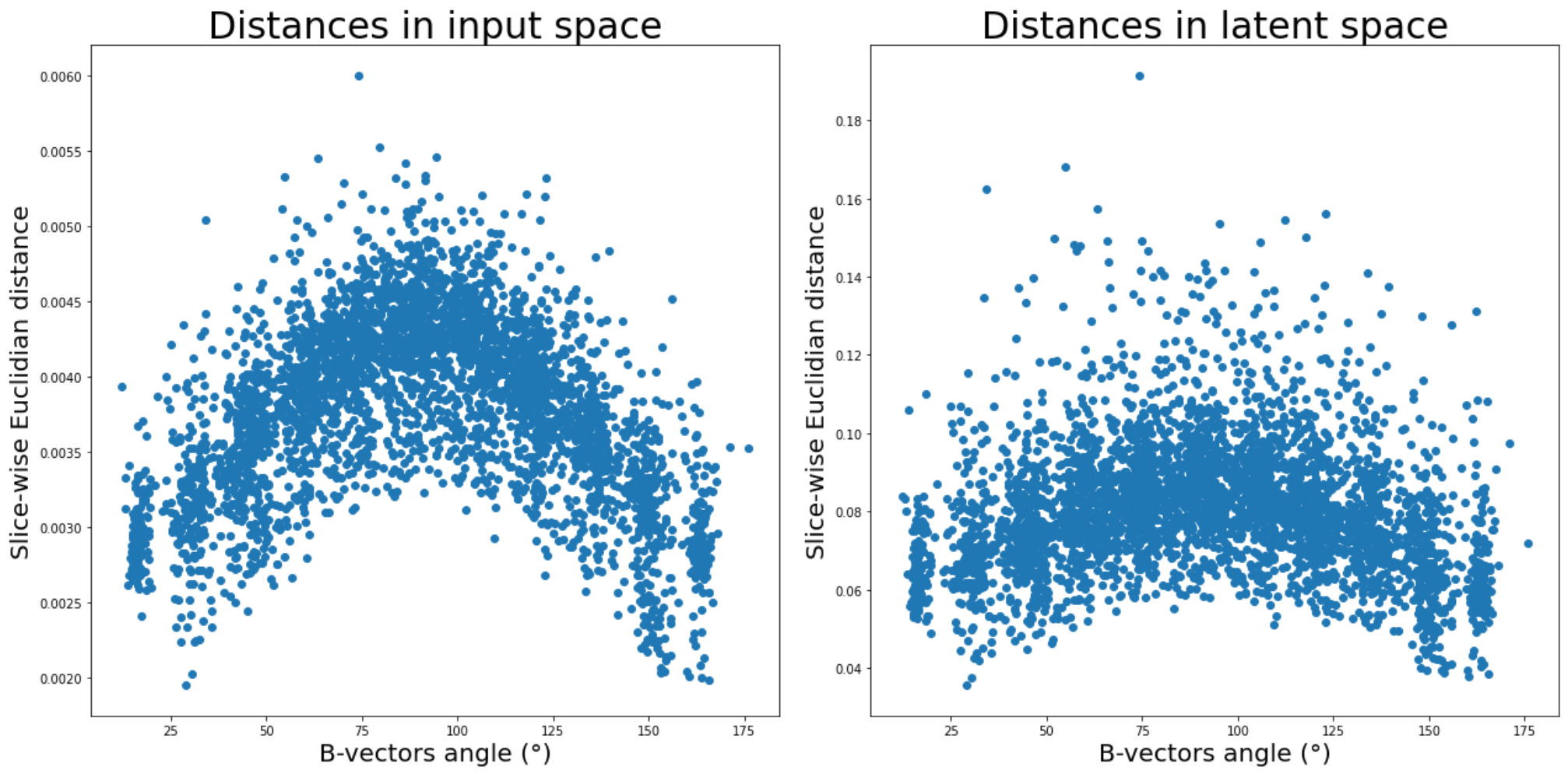
Average pair-wise slice distance between gradient direction volumes in input space (left) and latent space (right).

Moreover, our network that was exclusively trained on b0 images, was able to generalize to b1000. In fact the signal similarity between b0 and DW images was also used in Coupé et al. (2013) in an inverse problem paradigm in which a b0 prior was incorporated to reconstruct b700 volumes.

Figure 4 illustrates qualitative results and absolute errors for N=1 with respect to the GT (right) between the best interpolation baseline (trilinear, left) and the autoencoder enhancement (middle) for b1000. We overall saw from these representative examples, higher absolute intensities in the Linear-1 configuration than in the AE-1. However, ventricles are less visible when using autoencoder. We hypothesize this is because of their higher intensity in b0 images on which the network was trained.

**Figure 4.**
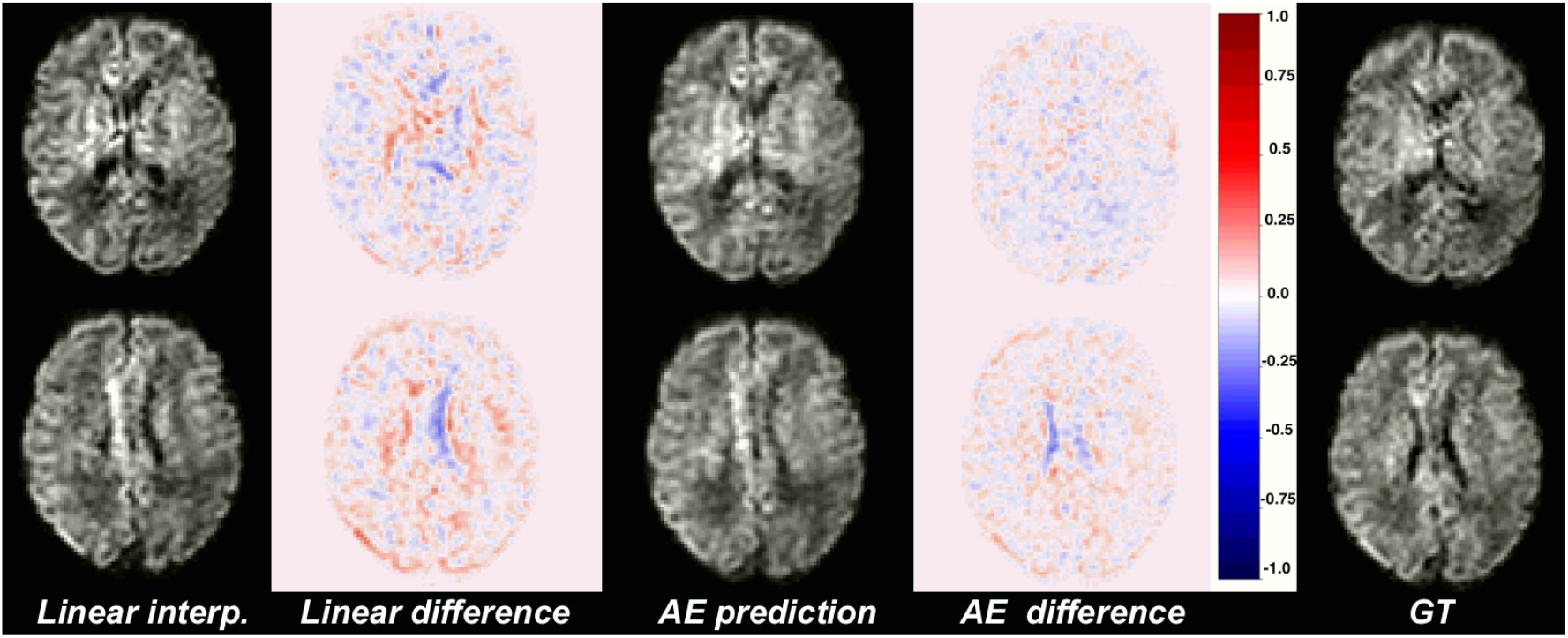
Illustration of the error difference in b1000 with respect to the ground truth (GT) for the best baseline method (trilinear, left) and autoencoder (AE, middle) enhancement.

The average MSE with respect to original DW-MRI signal within the whole brain is shown in Figure 5 for both the autoencoder enhanced volume and the baseline methods (trilinear, tricubic and B-spline), for the configurations where one (Method-1) or two (Method-2) slices were removed. The first observation was the expected higher error for the configuration where two slices are removed (N=2), independently of the method used. Additionally, the autoencoder enhancement clearly outperformed the baseline methods in all configurations (paired Wilcoxon signed-rank test p<1.24e-09). Particularly, the more slices we remove the higher the gap between the baseline interpolation methods and the autoencoder enhancement. For b0, the MSE gain was around 0.0005 for N=1 and 0.0015 for N=2 between the autoencoder and the average baseline method (Spline-1 v.s. AE-1 and Linear-2 v.s AE-2). For b1000, the gain between AE-1 and Cubic-1 was 0.0007, respectively 0.0015 between AE-2 and Cubic-2.

**Figure 5.**
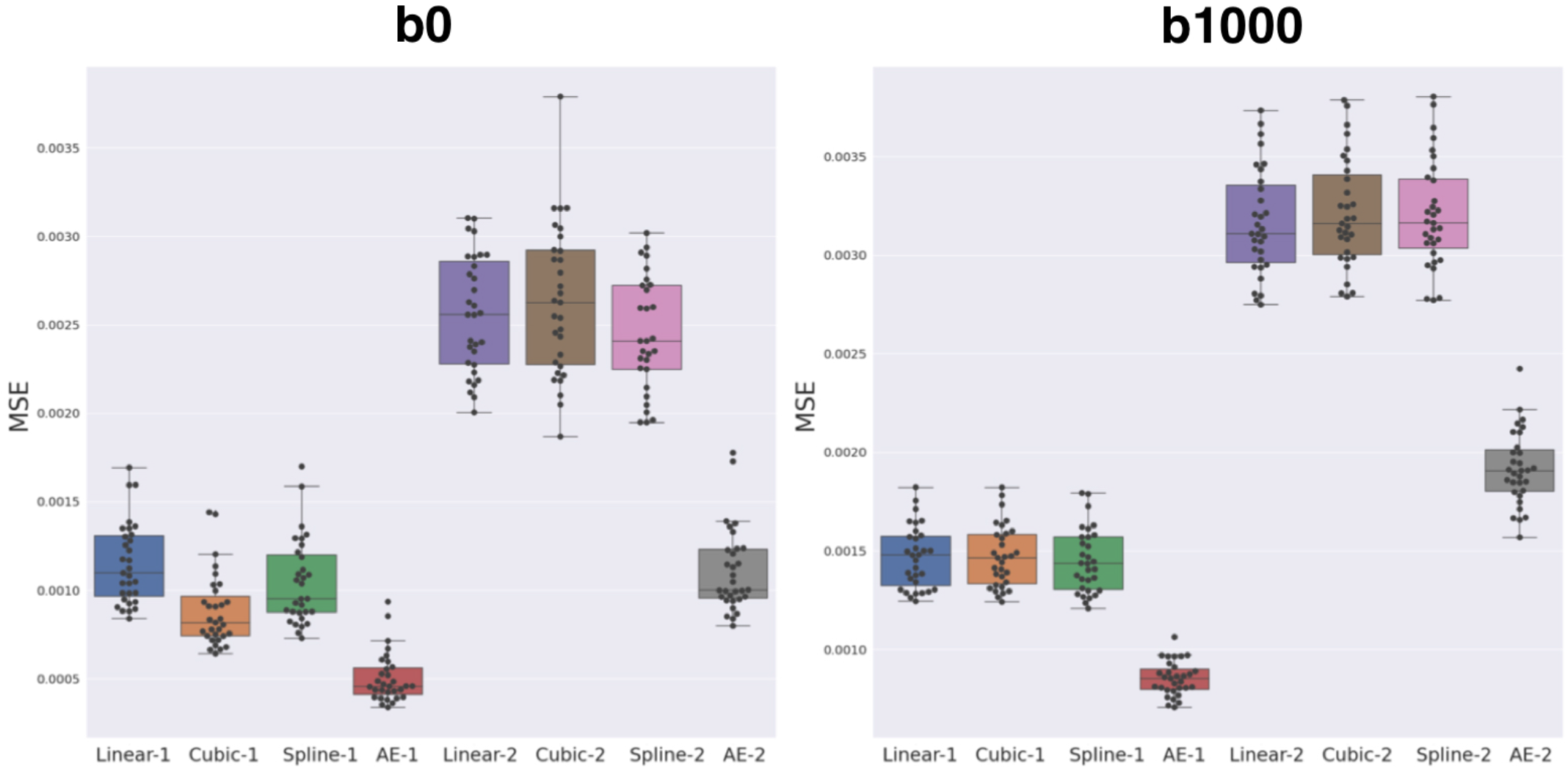
Mean squared error (MSE) between the three baseline methods (linear, cubic and B-spline 5*^th^* order) and autoencoder (AE) enhancement both for b0 (left) and b1000 (right). Two configurations were assessed: either N=1, i.e. removing one slice and interpolate/synthesize it (Linear-1, Cubic-1, Spline-1, AE-1) or N=2, i.e. the same approach with two slices (Linear-2, Cubic-2, Spline-2, AE-2). The autoencoder has a significantly lower MSE when compared to each respective best baseline method (paired Wilcoxon signed-rank test p<1.24e-09).

The overperformance of the autoencoder is also shown overall to the DTI maps, where MD, AD and RD were better approximated when compared to the best baseline method (linear interpolation), particularly in the configuration where two slices were removed (Figure 6). However, the FA showed the opposite trend especially for the configuration where one slice was removed (AE-1 v.s. Linear-1). However, FA for white matter-like structures (’WM’, corpus callosum and brainstem) showed higher performance with the autoencoder as depicted for each structure in Figure 7. In fact, by plotting colored FA for these two configurations we observed that the autoencoder generates tracts that were consistent with the GT. For instance, autoencoder enhancement showed higher frequency details around the superficial WM area (Figure 8, top row) and removed artefacts between the internal capsules better than the linear method (Figure 8, bottom row). However, in some cases, the baseline method better depicted tracts such as in the corpus callosum (Figure 8, middle row). Figure 9 shows similar comparisons for MD in different brain regions between the baseline method (Linear), the autoencoder and the GT. Overall, quantitatively, for structures in the case where two slices were removed, the autoencoder enhancement outperformed the best baseline method in 15 out of 16 configurations (Figure 7). However, it is not always the case when one slice is removed such as in the AD of the brainstem.

**Figure 6.**
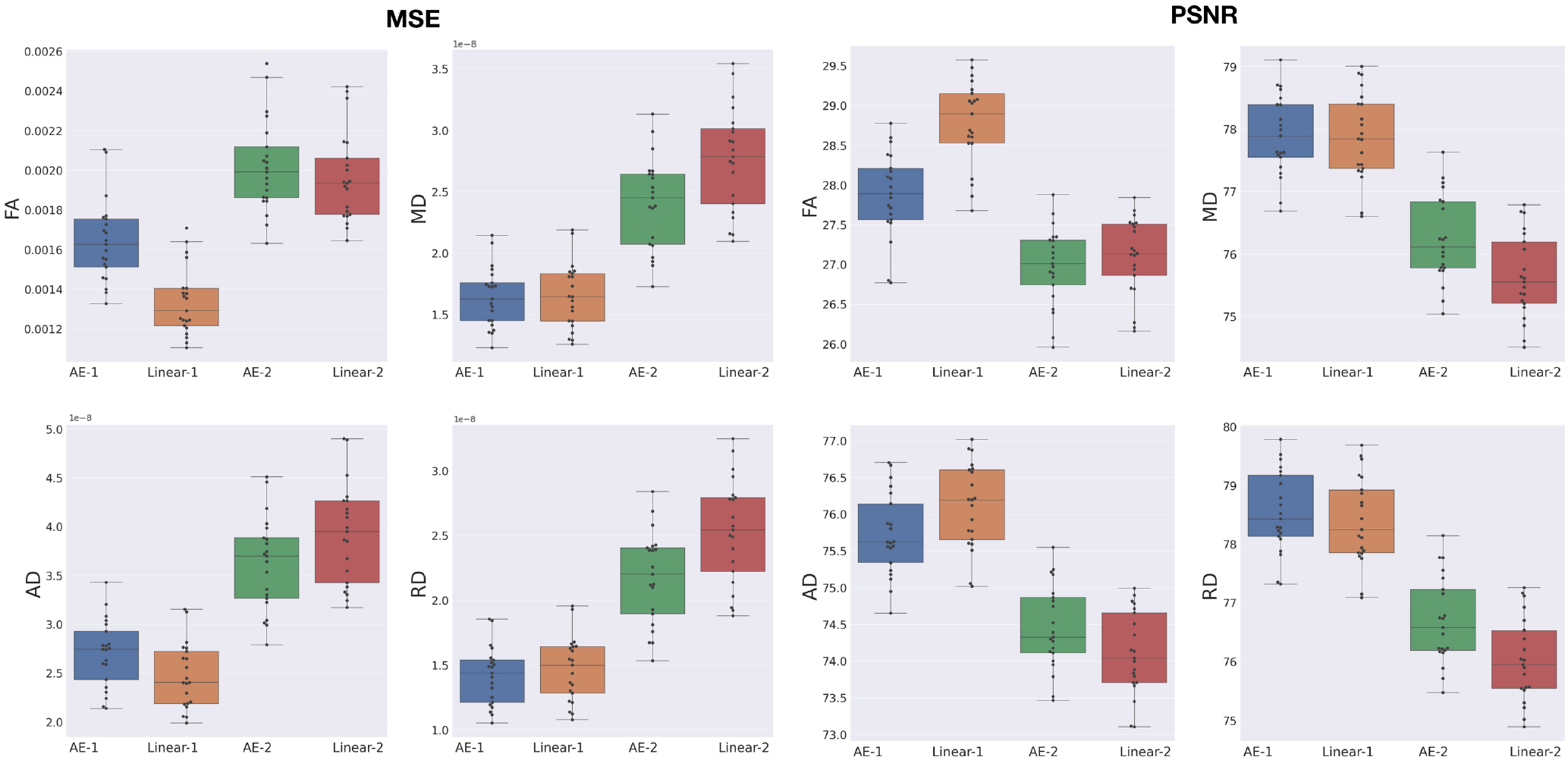
MSE and PSNR between the best baseline (Linear) and the autoencoder (AE) enhancement for whole-brain diffusion tensor maps, when removing and synthesizing/interpolating one or two slices.

**Figure 7.**
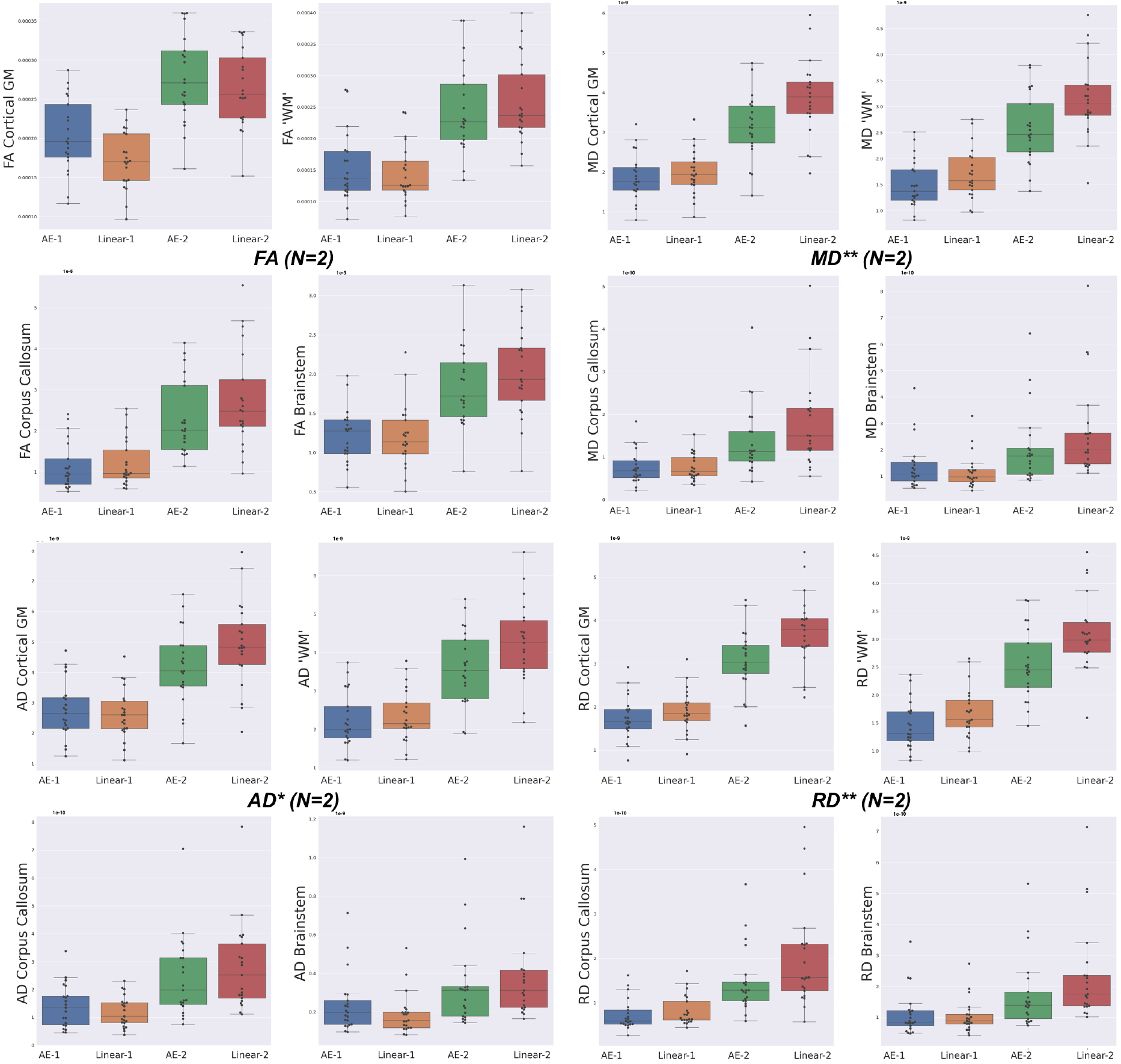
Mean squared error (MSE) with respect to the GT of the best baseline method (Linear) and the autoencoder (AE) enhancement in the different brain structures (Cortical Gray Matter (GM), White Matter (WM), Brainstem and Corpus Callosum) for each diffusion tensor map (FA, MD, AD and RD) for one slice removal (N=1) and two slices removal (N=2). Comparing the DTI maps of the merged brain region labels, we found that the AE-2 outperforms other conventional methods for MD, RD and AD. (Paired Wilcoxon signed-rank test: ** p<0.0073 - * p<0.051)

**Figure 8.**
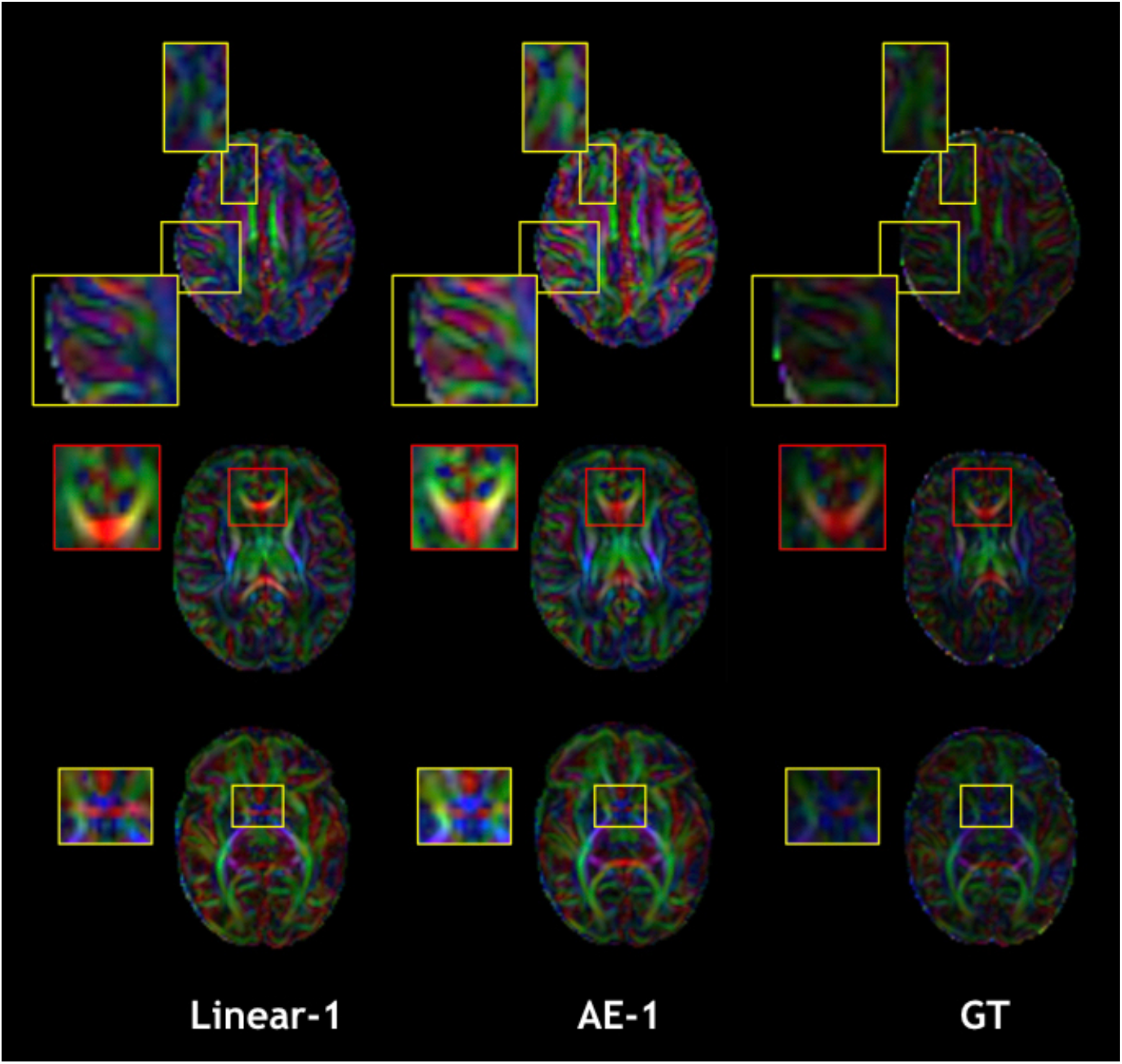
Qualitative comparison of colored FA in the one slice removed configuration for the best baseline interpolation method, i.e. Linear-1 (left), autoencoder enhancement AE-1 (middle) and GT (right). The red frame area highlight to a region where the linear interpolation shows a more accurate result.

**Figure 9.**
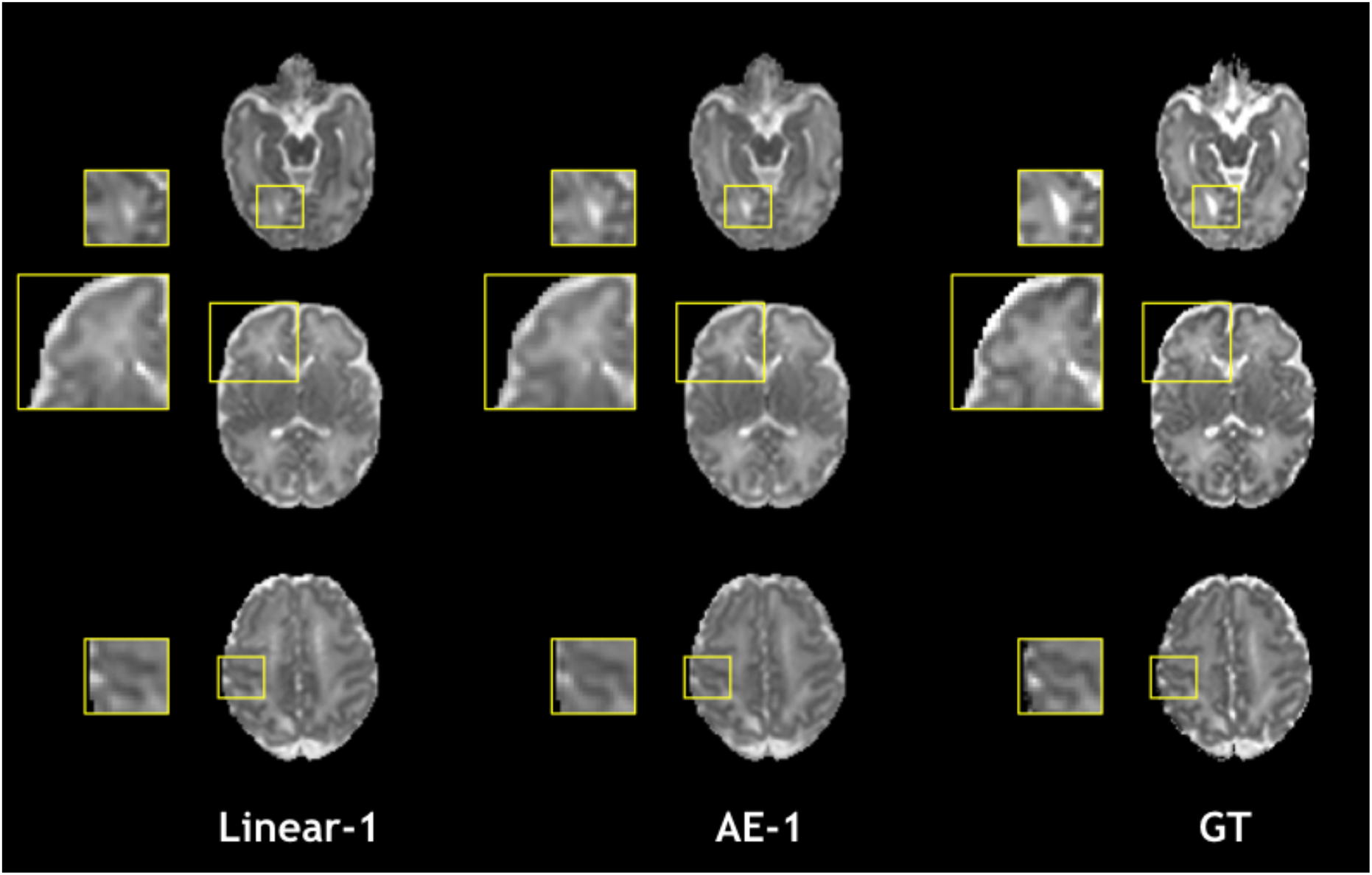
Qualitative comparison of mean diffusivity (MD) in the one slice removed configuration for the best baseline interpolation method, i.e. Linear-1 (left), autoencoder enhancement AE-1 (middle) and GT (right).

Figure 10 shows how our autoencoder was robust to reasonable amounts of noise. In fact, simply encoding and decoding the noisy input generates a slice that was closer to the GT than the noisy slice, as depicted for different levels of noise for both b0 and b1000.

**Figure 10.**
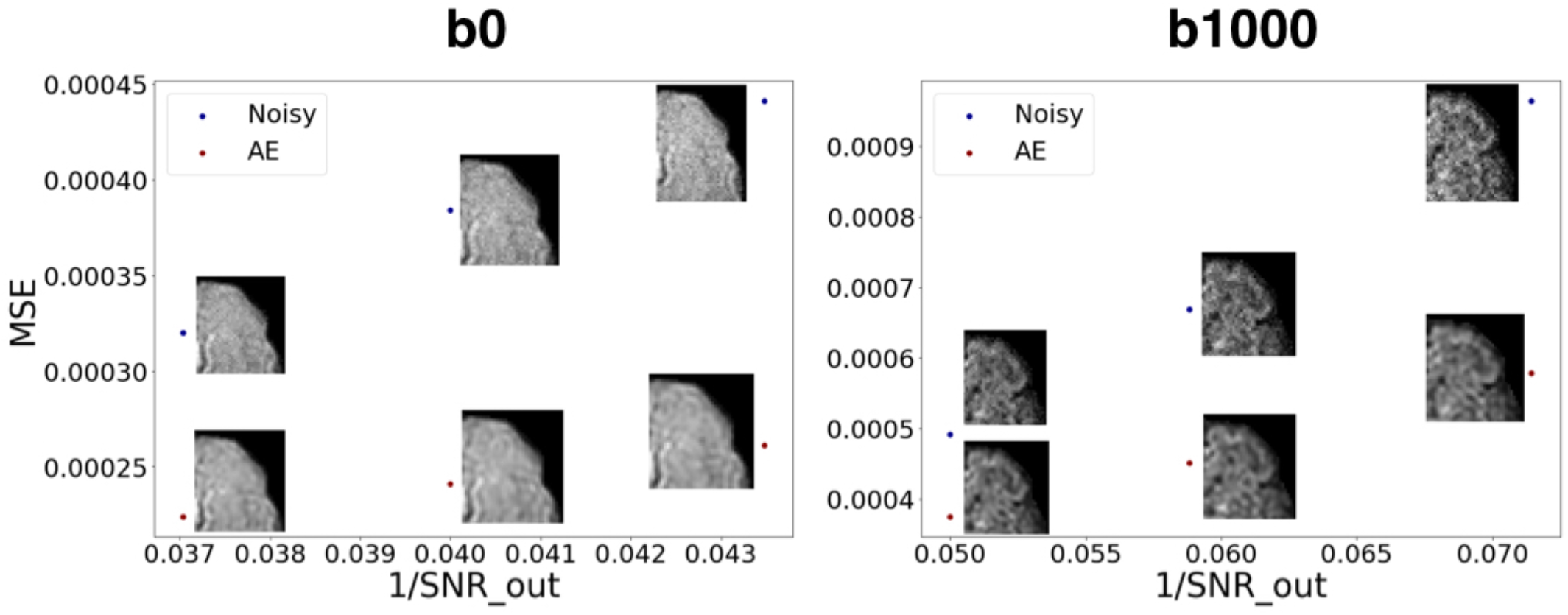
MSE between noisy images and the GT v.s. encoded-decoded noisy images and the GT. SNR out is the desired SNR of the output in the Rician noise formula (Subsection 2.3.1). We notice the robustness of the autoencoder to growing levels of noise both for b0 images (left) and b1000 images (right).

### 3.2 Fetuses

Figure 11 illustrates inter-volume motion between five diffusion-weighted volumes where we notice a severe signal drop in the seventh direction (discarded volume).

**Figure 11.**
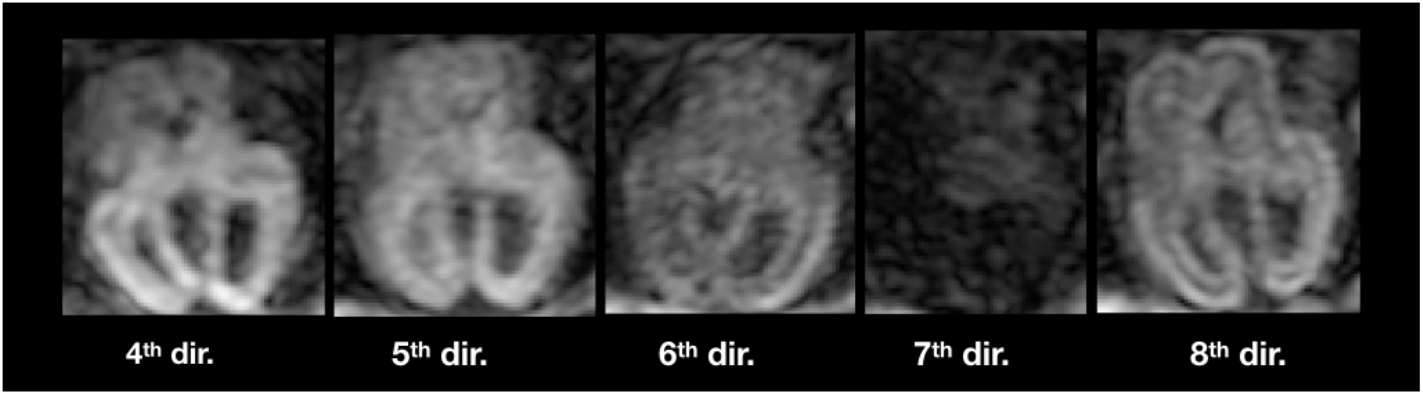
Illustration of inter-volume motion in five different gradient directions. Note the severe signal drop in the seventh direction because of motion.

The autoencoder trained on pre-term b0 images was able to coherently enhance fetal acquisitions both at b0 and DW-MRI volumes at b700. The network was able to learn low-level features that could generalize over anatomy, contrast and b-values. Corresponding FA and colored FA for a still subject (35 GW) are illustrated in Figure 12 (top) where we clearly see the coherence of the two synthesized images as we go from one original slice to the next one. In fact, both the corpus callosum and the internal/external capsules follow a smooth transition between the two slices. Similarly, Figure 12 (bottom) exhibits MD and FA for a moving subject (24 GW) where we also notice, particularly for the MD, the smooth transition between the originally adjacent slices. FA and MD for the remaining subjects are shown in Figure S1 in Supplementary Material.

**Figure 12.**
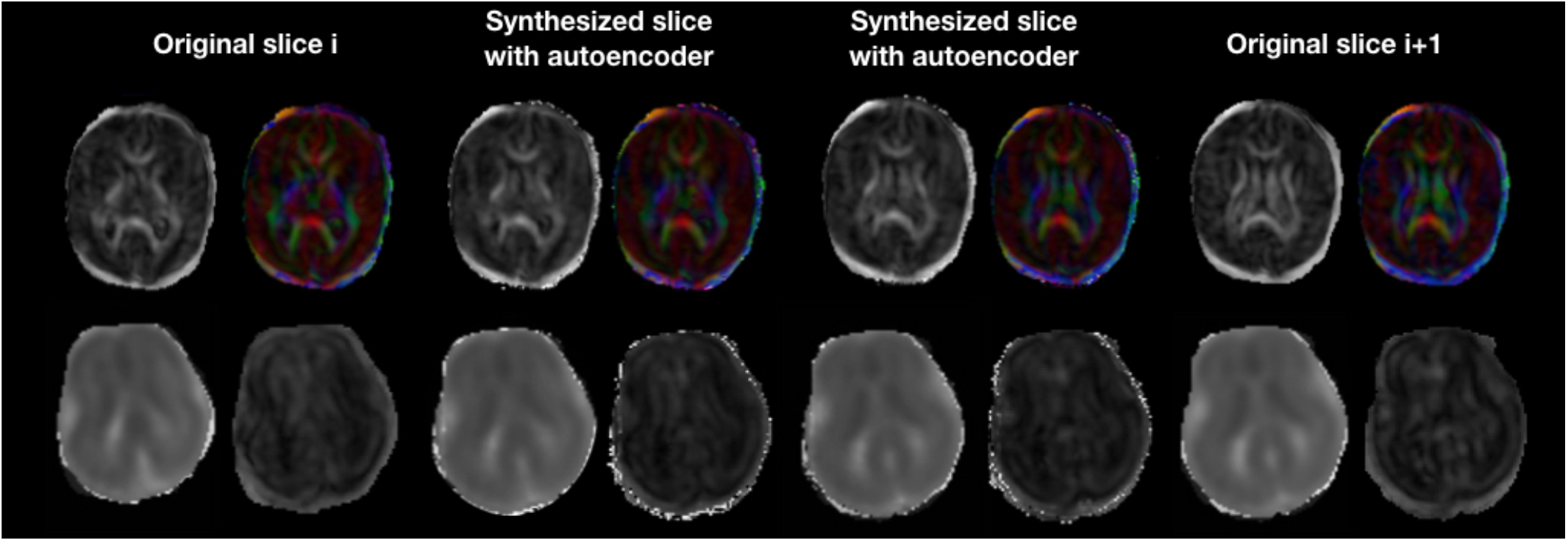
Colored FA and FA (top row) illustration of autoencoder enhancement between two original adjacent fetal slices in a still subject (35 GW). The bottom row shows a similar illustration of MD and FA for a moving subject (24 GW).

The splenium and genu of corpus callosum were only sufficiently visible in the three late GW subjects (27, 29 and 35 GW) subjects. Figure 13 shows quantitative results for FA and MD in the two structures. Both maps fall into the range of reported values in the literature (Wilson et al., 2021) for the respective gestational age, for original and autoencoder enhanced volumes.

**Figure 13.**
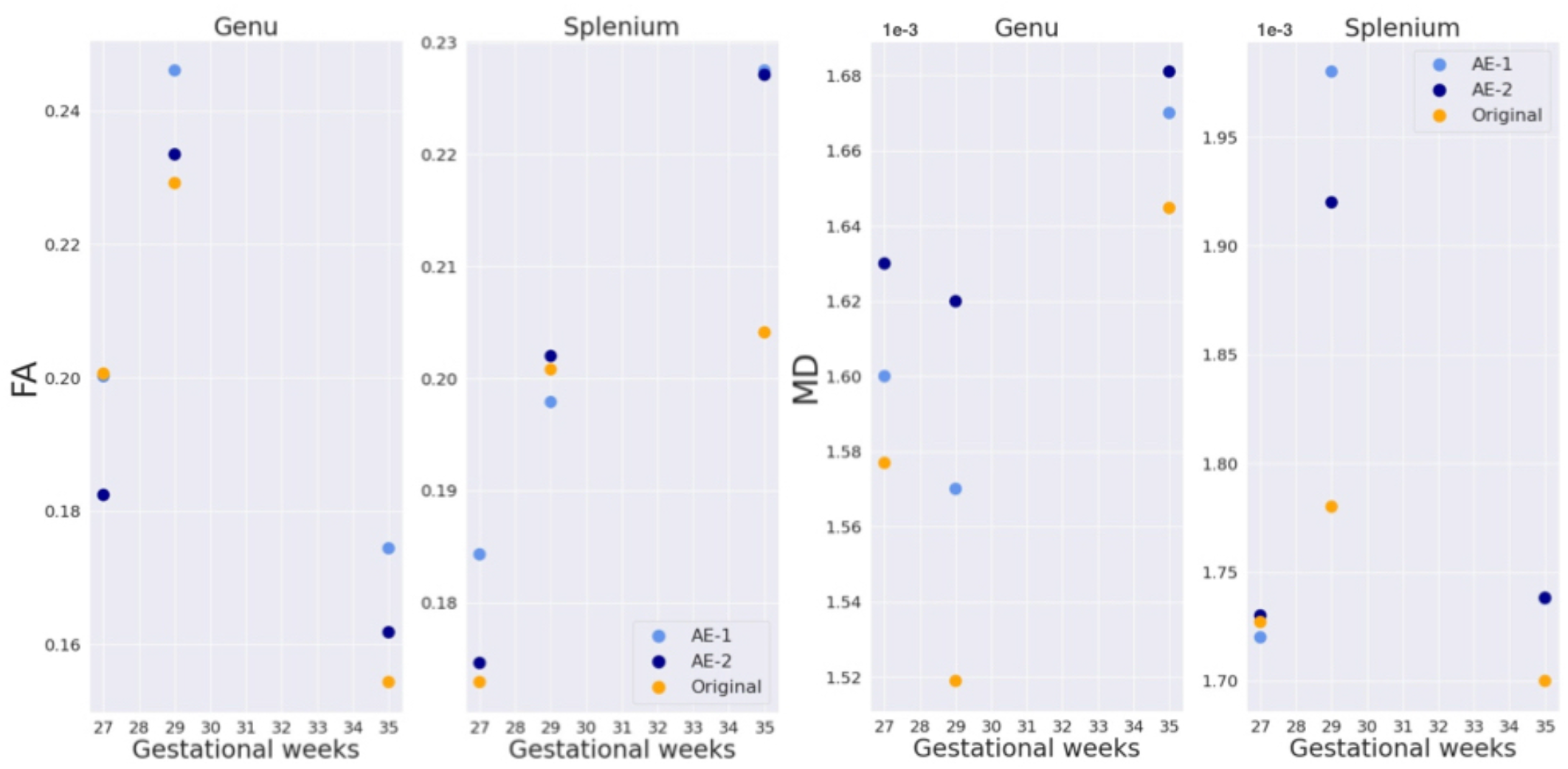
FA and MD in genu and splenium of corpus callosum for three subjects (27, 29 and 35 GW)

## 4 DISCUSSION

In this work we have shown that (1) autoencoders can be used for through-plane super-resolution in diffusion MRI, (2) training on b0 images can generalize to gradient diffusion volumes of different contrasts and (3) as a proof of concept, training on pre-term anatomy can generalize to fetal images.

In fact, we have demonstrated how autoencoders can realistically enhance the resolution of DW-MRI perinatal brain images. We have compared it to conventionally used methods such as trilinear, tricubic and B-spline interpolations both qualitatively and quantitatively for pre-term newborns of the dHCP database. Resolution enhancement was performed at the diffusion signal level and the downstream benefits propagated to the DTI maps.

Additionally, our network that was solely trained on non-diffusion weighted images (b0) was able to generalize to a b1000 contrast. In fact, the most intuitive approach is to infer b1000 images using a network trained on b1000. We have indeed tried but the network did not converge for the majority of the folds. This might be due to the high variability of b1000 images across directions and their inherently low SNR. However, in the one fold that the network converged, it slightly underperformed the network that was trained on b0 only, on both b1000 pre-term and b700 fetal images. Moreover, being b-value independent is a desirable property since different b-values are used in different centers, in particular for clinical fetal imaging (400, 500, 600, 700 *s/mm*^2^) (Fogtmann et al., 2013; Jakab et al., 2015, 2017; Marami et al., 2017; Deprez et al., 2019). In fact, the same b0 network trained on pre-term data was generalized to b700 fetal images where we qualitatively show its advantage, hence supporting the utility of pre-term data for fetal imaging such as in Karimi et al. (2021), where they have used pre-term colored FA and DW-MRI fetal scans to successfully predict fetal colored FA using a convolutional neural network. Furthermore, FA and MD of the corpus callosum which were generated using the autoencoder enhanced volumes are in the range of values provided by a recent work (Wilson et al., 2021). This a necessary but non sufficient condition for the validity of our framework in fetal data.

Notably, our trained network was able to reduce the noise from the data by learning the main features across images for different noise levels. This can be explained by two points. First, our autoencoder was exposed to different low levels of noise (as the dHCP data was already denoised) and hence the encoded features of the latent space are ought to be noise independent. Secondly, generative autoencoders intrinsically yield high SNR outputs due to the desired smoothness property of the latent space (Berthelot et al., 2018).

The proposed framework could be applied to correct for anisotropic voxel sizes and can be used for slice outliers recovery in case of extreme motion artefacts for example. In fact, the artificially removed middle slices in our experiments can represent corrupted slices that may need to be discarded or replaced using interpolation (Niethammer et al., 2007; Chang et al., 2005; Andersson et al., 2016). Our autoencoder can hence be used to recover these damaged slices using neighboring ones.

The power of our method compared to conventional interpolations resides in two points. Firstly, the amount of data used to predict/interpolate the middle slice. While only two slices will be used in traditional interpolation approaches, our method will in addition take advantage of the thousands of slices to which the network has been exposed and from which the important features have been learned (without any supervision) in the training phase. Secondly, based on the manifold hypothesis, our method performs interpolations in the learned encoding space, which is closer to the intrinsic dimensionality of the data (Chollet, 2017), and hence all samples from that space will be closer to the true distribution of the data compared to a naive interpolation in the pixel/voxel space.

Although our network performed quantitatively better than conventional interpolation methods in preterm subjects, its output is usually smoother and hence exhibits less details. This is a well-known limitation of generative autoencoders, such as variational autoencoders, and the consequence of the desirable property of making the latent space smooth (Berthelot et al., 2018). Generative Adversarial Networks (Goodfellow et al., 2014) can be an interesting alternative to overcome this issue. However, they have other drawbacks as being more unstable and less straightforward to train (Mescheder et al., 2018) than autoencoders.

In this work, qualitative results only were provided on fetal DW-MRI. We are limited by the lack of ground truth in this domain, hence our results are a proof of concept. The future release of the fetal dHCP dataset will be very valuable to further develop our framework and proceed to its quantitative assessment for fetal DW-MRI.

In future work, we want to add random Rician noise in the training phase to increase the network robustness and predictive power. We also want to extend the autoencoding to the angular domain by using spherical harmonics decomposition for each 4D voxel, and hence enhancing both spatial and angular resolutions (Ma and Cui, 2021).

Although *unsupervised* learning via autoencoders has been recently used in DW-MRI to cluster individuals based on their microstructural properties (Chamberland et al., 2021; Rokem, 2021)), this is to the best of our knowledge, the first *unsupervised* learning study for super-resolution enhancement in DW-MRI using autoencoders.

As diffusion fetal imaging suffers from low through-plane resolution, super-resolution using autoencoders is an appealing method to artificially but realistically overcome this caveat. This can help depict more precise diffusion properties through different models such as DTI or ODFs and potentially increase the detectability of fiber tracts that are relevant for the assessment of certain neurodevelopmental disorders (Jakab et al., 2017).

## 5 ADDITIONAL REQUIREMENTS

### CONFLICT OF INTEREST STATEMENT

The authors declare that the research was conducted in the absence of any commercial or financial relationships that could be construed as a potential conflict of interest.

### AUTHOR CONTRIBUTIONS

H.K.: Original idea, conceptualized the research project, performed the technical analysis, wrote the manuscript and integrated all revisions. M.B.C.: Conceptualized, designed and supervised the research project, contributed to the manuscript and to the final revision, and provided funding. E.C.R.: Contributed to the conceptualization of the research project and revised the manuscript. H.L.: Helped in the data generation and revised the manuscript. P.Du.: Helped in the technical analysis and revised the manuscript. G.G.: Helped in the preprocessing of the fetal data and revised the manuscript. Y.A.: Helped in the preprocessing of the fetal data and acknowledged the manuscript. M.K.: Helped in the processing of the fetal data and acknowledged the manuscript. A.J.: Provided the fetal data and revised the manuscript.

### FUNDING

This work was supported by the Swiss National Science Foundation (project 205321-182602, grant No 185897: NCCR-SYNAPSY- ”The synaptic bases of mental diseases” and the Ambizione grant PZ00P2_185814).

## ACKNOWLEDGMENTS

This work was supported by the Centre d’Imagerie BioMédicale (CIBM) of the University of Lausanne (UNIL), the Swiss Federal Institute of Technology Lausanne (EPFL), the University of Geneva (UniGe), the Centre Hospitalier Universitaire Vaudois (CHUV), the Hôpitaux Universitaires de Genève (HUG), and the Leenaards and Jeantet Foundations. We also thank Athena Taymourtash for the help in the acceleration of the technical analysis and Samuel Lamon (MD) for the help in the segmentation of the corpus callosum.

## DATA AVAILABILITY STATEMENT

The pre-term newborns dataset analyzed in this study is a subset of the developing Human Connectome Project (dHCP) dataset. This dataset is publicly available at: http://www.developingconnectome.org/data-release/data-release-user-guide/.

The fetal data is from the University Children’s Hospital Zürich (KISPI) and cannot be shared with the public at this time.

## Supplementary Material

### 1 SUPPLEMENTARY TABLES AND FIGURES

**Table S1.** Main settings of acquisition protocols in fetal brain diffusion MRI.

|  | Field strength & brand | #Directions | B-values (s/mm<sup>2</sup>) | Orthogonal stacks | Resolution (mm<sup>3</sup>) | Acquisition time in minutes:seconds | Main sequence parameters (TE/TR in ms) | Year |
| --- | --- | --- | --- | --- | --- | --- | --- | --- |
| Jakab et al., Neuroimage: Clinical 2017 | 1.5T/3T GE | 15 | 700 | 3 axial and 1-2 coronal | 1x1x3-5 | 1:21 per 4D volume | TE = min(65)<br>TR = 2200 | 2020 |
| Deprez et al., IEEE Transactions on Medical Imaging 2019 | 3T Phillips | 32 | 750 | ? | 2x2x3.5 | ~5:00 | TE = 75<br>TR = 7500 | 2020 |
| Christiaens et al., ISMRM 2019 (DHCP) | 3T Phillips | 80 | 400<br>1000 | ? | 2x2x2 | ~15:00 (whole dataset) | TE = 75<br>TR = 6100 | 2019 |
| Marami et al., Neuroimage 2017 | 3T Siemens | 12 | 500 | 2-8 axial/<br>coronal | 2x2x2-4 | 0:50-1:30 per 4D volume | TE = 60<br>TR = 3000/4000 | 2017 |
| Jakab et al., Neuroimage 2015 | 1.5T Phillips | 15 | 700 | Axial | 0.94x0.94x3.3 | ? | TE = 90<br>TR = 1745 | 2015 |
| Fogtmann et al., IEEE Transactions on Medical Imaging 2014 | 1.5T Siemens | 10 | 500<br>600<br>700 | 2 axial<br>2 sagittal<br>0-2 coronal | ~ 2x2x2 | 00:10 per 4D volume | TE = 96<br>TR = 4000-6000 | 2014 |

**Table S2.**
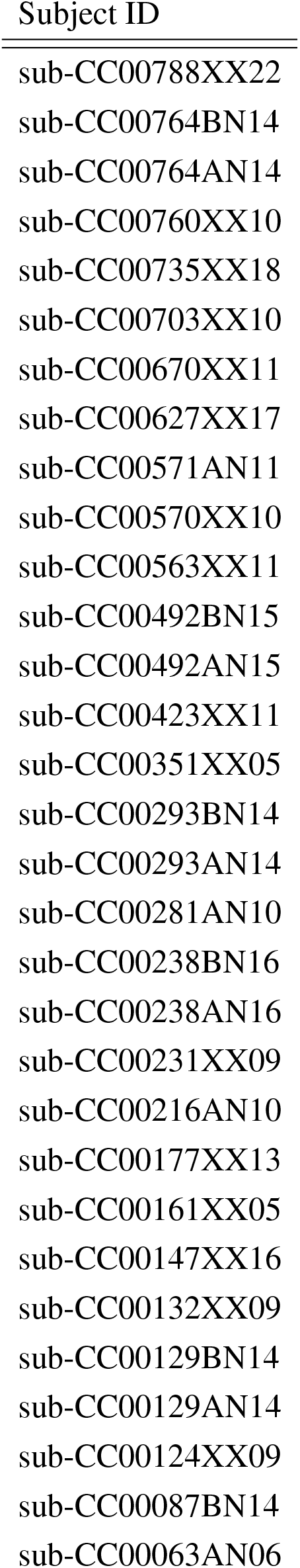
IDs of the pre-term subjects used from the developing Human Connectome Project (dHCP) dataset.

**Table S3.** Discarded volumes per each 4D acquisition for three subjects. All the volumes were kept for the remaining subjects.

|  | Axial-1 | Axial-2 | Axial-3 |
| --- | --- | --- | --- |
| Motion subject 1 | - | Vol 6,7,15 | Vol 11 |
| Motion subject 2 | Vol 2,3,4,5,6,7 | Vol 11,14 | Vol 14 |
| Still subject 1 | All volumes except vol 0 (b0) | - | All volumes except vol 0 (b0) |

**Figure S1:**
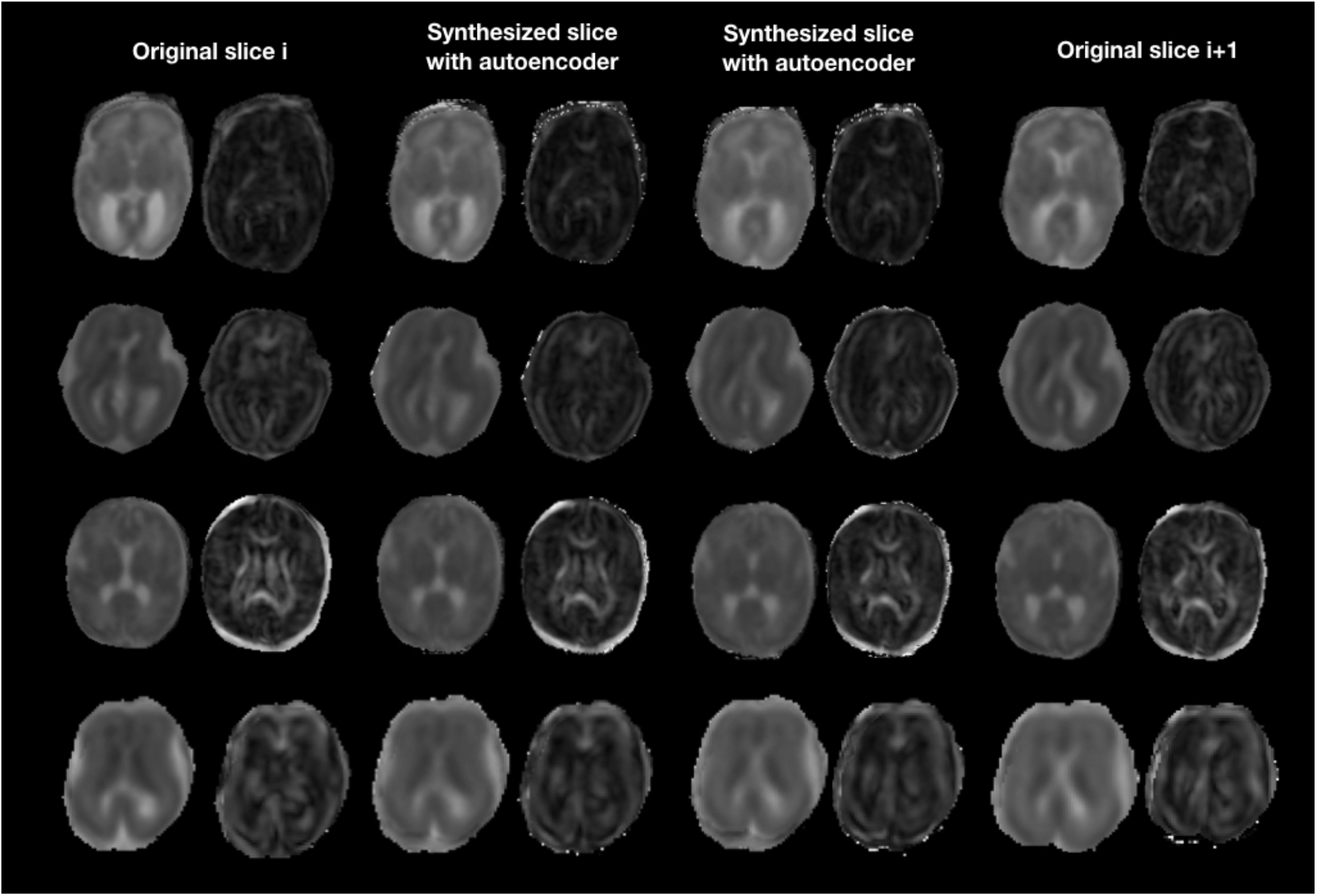
MD and FA illustration of AE enhancement between two adjacent fetal slices in two moving subject (top rows: 27, 23 GW) and two still subjects (bottom rows: 35, 24 GW).

## Footnotes

1 www.github.com/Medical-Image-Analysis-Laboratory/Perinatal_SR_Autoencoder

## Notes

### Competing Interest Statement

The authors have declared no competing interest.

